# Spontaneous emergence of topographic organization in a multistream convolutional neural network

**DOI:** 10.64898/2026.02.23.707577

**Authors:** Hiroshi Tamura

## Abstract

Neurons in the cerebral cortex are organized topographically. In the primate visual cortex, neighboring neurons often respond to similar stimulus parameters, such as receptive field position, orientation, color, and spatial frequency. Preferred stimulus parameters change smoothly across the cortical surface. If such topographic organization plays an important role in computation, it is likely to emerge in artificial neural networks. In this study, a multistream convolutional neural network was constructed in which filters in the first convolutional layer were arranged in a two-dimensional filter matrix according to their output connections. The network was trained using supervised learning for image classification. Although adjacent filters in the filter matrix can develop any structure in principle, they acquire similar degrees of orientation and color selectivity. Moreover, they prefer similar orientations, hues, and spatial frequency. The similarity decreases with distance between filters in the matrix. Furthermore, neural-network model instances that have a strong relationship between filter distance and filter-property similarity performed better than those with a weak relationship. These results suggest that topographic organization emerges spontaneously in an artificial neural network and plays an important role in model performance, suggesting the importance of topographic organization for computations performed by artificial and biological neural networks.

## Introduction

Topographic organization is observed in a variety of stimulus parameters in the primary visual cortex (V1) of primates. In V1, neurons respond to stimuli presented at a specific location in the visual field, which is called the receptive field (RF). Neighboring neurons have similar RF positions and RF areas, which change smoothly across the cortical surface^1-3^. Neighboring neurons in the primary visual cortex also encode similar types of visual submodality. Color-selective neurons are more abundant in cytochrome oxidase blobs than in inter-blob regions, whereas orientation-selective neurons are more abundant in inter-blob regions than in blob regions^4-10^. Furthermore, neighboring neurons tend to be activated by the same stimulus parameter. Moreover, this stimulus parameter changes smoothly across the cortex and is organized topographically. For example, the preferred orientation of neighboring V1 neurons is similar and changes smoothly across the cortex^11-14^. The preferred color and spatial frequency of a V1 neuron are also similar to those of neighboring neurons and change smoothly across the cortex^15-18^.

One possible reason for the ubiquity of topographic organizations in the cerebral cortex concerns anatomical constraints. Because neighboring neurons are likely to receive inputs from the same axons^19^, they are likely to share response properties. The degree of input sharing decreases gradually across the cortex and results in topographic organization.

Another possible reason for the emergence of topographic organizations involves computational requirements. In the primary visual cortex, computation is primarily performed among neurons encoding similar parameters ^20, 21^ to enhance spatial contrast, improve stimulus tuning, and provide invariant response properties. If neurons are organized in a topographic manner, total axon length, which is proportional to metabolic resources and computational time, can be minimized. However, the computational importance of topographic organization in the brain remains unclear^22-24^.

Recently, convolutional neural networks (CNNs) have been used to investigate the computational significance of functional organization in the brain^25^. CNNs are hierarchically organized feed-forward networks that consist of multiple sets of layers. Each set of layers performs convolution, thresholding, and pooling^26^. The filter weights for convolution are initially set to random values and modified during training. Recently, functional module-like organization was found to spontaneously emerge in multistream CNNs^27, 28^.

In the present study, to examine the computational significance of topographic organization, a multistream CNN was constructed such that filters in the first convolutional layer (conv1) are placed in a two-dimensional (2D) filter matrix according to the output connections. The aim was to examine whether filters in conv1 are arranged in a topographic manner in a 2D filter matrix. The underlying rationale is as follows: if the properties of conv1 filters change smoothly in the filter matrix, then the presence of topographic organization in the filter matrix of conv1 is required for information processing. In a previous study^29^, topographic organization emerged in an artificial neural network when filters were placed retinotopically and the spatial loss, which constrains filter similarity between adjacent filters, was introduced. In the present study, because no constraints were placed on the input weight structure for the conv1 filters nor the similarity of the filter properties between adjacent filters in the filter matrix, the models were—in principle—able to develop any filter structure. Nonetheless, adjacent filters in the filter matrix developed similar properties, and similarity changed gradually across the filter matrices. Furthermore, a neural-network model instance that had a strong relationship between filter distance and filter-property similarity outperformed one with a weak relationship. These results suggest that topographic organization in the filter matrices of conv1 filters is required to obtain the optimal task performance of a multistream CNN.

## Results

A multistream CNN (tmcAlexNet; Fig. 1) was constructed based on a multistream CNN with convergence (mcAlexNet)^28^. tmcAlexNet consists of five hierarchically organized convolutional layers (conv1–conv5) and three pooling layers (Max-pool). Conv1 contains 256 filters (0–255), and conv2 contains 256 streams (s1–s256), each equipped with 12 filters. Each conv2 filter receives converging inputs that come from four conv1 filters. For example, the 12 filters in s1 of conv2 receive converging inputs from filters 0, 1, 2, and 3 of conv1 (Fig. 1, red lines), and the 12 filters in s2 of conv2 receive converging inputs from filters 1, 3, 4, and 6 of conv1 (Fig. 1, green lines), and so on. As a result, filter 3 in conv1 shares target filters in conv2 with eight other conv1 filters (filters 0–2, 4, 6, 8, 9, and 12). Similarly, filter 6 shares target filters in conv2 with eight other conv1 filters (filters 1, 3–5, 7, 9, 12, and 13).

**Figure 1.**
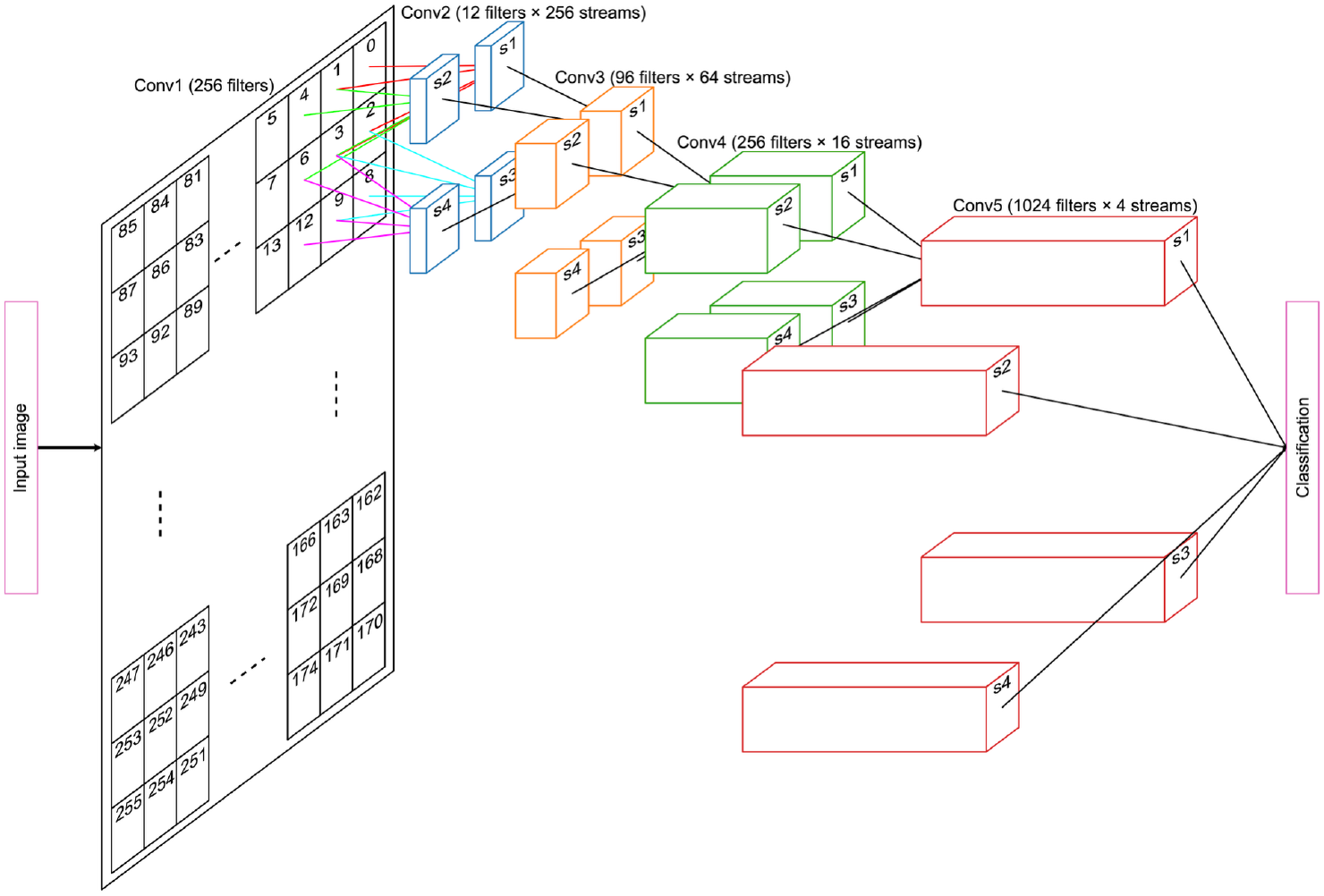
Architecture of a multistream convolutional neural network (tmcAlexNet). The numbers of filters and streams of the convolutional layers (conv1-5) are indicated in parentheses. Only a part of the filters in conv1 and some of the streams for conv2-4 are shown for visualization purposes. Each conv1 filter is placed on a 2D matrix (filter matrix) according to the output connections. Note that the placement of conv1 filters is based on the degree of sharing in the target conv2 filters and is not related to retinotopic position.

Based on the degree of sharing in the target filters, each conv1 filter is placed on a 2D matrix (filter matrix; Fig. 2A right, Supplementary Figure 1). For example, the eight conv1 filters (filters 0–2, 4, 6, 8, 9, and 12) that share target filters with filter 3 are placed around filter 3 in the filter matrix, and the eight filters (filters 1, 3–5, 7, 9, 12, and 13) that share target filters with filter 6 are placed around filter 6 in the filter matrix. This filter arrangement is inspired by the anatomical organization of the primary visual cortex, in which outputs from neighboring neurons tend to converge onto common targets^4, 30^.

**Figure 2.**
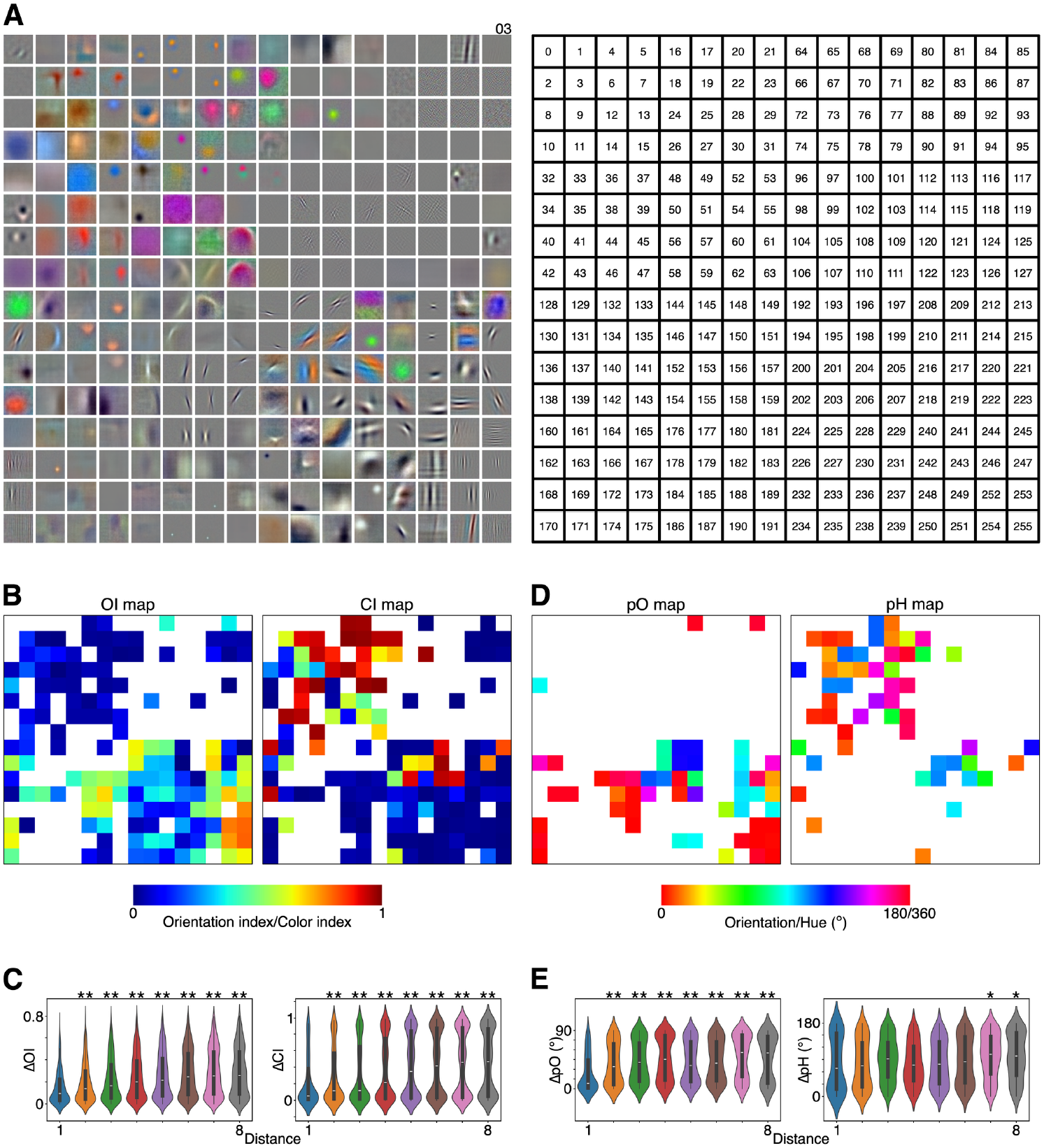
Spatial arrangement of filters in the filter matrix of the first convolutional layer (conv1) of tmcAlexNet for a representative model instance. ***A***, Visualization of the filter weights for conv1 filters (left). Filters are placed according to the filter matrix (right). The minimum and maximum weight values are scaled between 0 and 255 for visualization. ***B***, Maps of orientation index (OI map, left) and color index (CI map, right) for responsive filters (*n* = 143). Index values are color coded. ***C***, Comparisons of absolute differences in the OI (ΔOI, left) and CI (ΔCI, right). ***D***, Maps of the preferred orientation of orientation-selective filters (pO map; *n* = 53, left) and preferred hue of color-selective filters (pH map; *n* = 53, right). Preferred parameters are color coded. ***E***, Comparisons of absolute differences in preferred orientation (ΔpO, left) and preferred hue (ΔpH, right) across filter-distance groups. The violin plots show estimated kernel density. The white horizontal bar indicates the median or second quartile, and the bottom and top of the box indicate the first and third quartiles, respectively. Whiskers extend to data points that lie within 1.5 times of the inter-quartile range. Double and single asterisks indicate significant differences between the 1st and corresponding filter-distance group with *p* < 0.01 and *p* < 0.05, respectively.

Conv3 contains 64 streams (s1–s64), each equipped with 96 filters. Each conv3 filter receives converging inputs that come exclusively from the 48 filters in the four streams of conv2. Conv4 contains 16 streams (s1–s16), each equipped with 256 filters. Each conv4 filter receives converging inputs that come exclusively from the 384 filters in the four streams of conv3. Finally, conv5 contains four streams (s1–s4), each equipped with 1024 filters. Each conv5 filter receives converging inputs that come exclusively from the 1024 filters in the four conv4 streams. Outputs from the four conv5 streams are concatenated and fed into fully connected layers (FCs), and then passed to the output layer for classification. Note that the inputs to the conv1 filters are not constrained, and hence they can acquire any structure in principle.

tmcAlexNet was trained to classify 1,000 object categories using the ImageNet database^31^. Sixteen model instances of tmcAlexNet were trained for this study using randomly initialized parameters. After training, the top-5 accuracy of tmcAlexNet ranged from 0.479 to 0.495 on the validation set. This performance is lower than that of the original AlexNet^26^ but better than that of our previous study using mcAlexNet ^28^.

### Similarity in filter properties decreases with distance in the filter matrix of tmcAlexNet

After training, the conv1 filters in the tmcAlexNet model acquired a variety of filter structures (Fig. 2A). For example, filter 155 has an orientation index (OI)^27^ of 0.56, indicating that it is orientation selective, and its preferred orientation is 12.9°. However, it has a color index (CI) ^26^ of 0.025, indicating that the color selectivity of this filter is low. By contrast, filter 47 is color selective (CI, 0.98) but not orientation selective (OI, 0.028). The preferred hue of this filter is 2.3°.

A visual inspection of Figure 2A reveals that adjacent filters in the filter matrix tend to have similar properties. For example, the filters around orientation-selective filter 155 are also orientation selective, whereas the filters around color-selective filter 47 are also color selective, suggesting clustering based on the visual submodality in the filter matrix of conv1 of tmcAlexNet. This tendency is revealed by visualizing the OI and CI values on the filter matrix (OI and CI maps; Fig. 2B). Filters with high OIs are observed in the lower half of the region, whereas filters with high CI are observed in the top-left region. Thus, high OI filters are segregated from high CI filters. Indeed, there is a negative correlation between the OI and CI of responsive filters (*r* = -0.613, *p* = 3.61 × 10^-16^, *n* = 143, Spearman’s rank correlation).

To quantitatively examine the topographic organization of filter properties in the filter matrix, the relationship between the similarities in filter properties and filter distance is examined. The filter distance is defined using the following rule: the distance between a filter and the eight surrounding filters in the filter matrix is defined as one, and that between the center filter and those outside of the eight surrounding filters is two. In this way, the distance between the filters takes values from one to eight (Supplementary Figures 2 and 3) and each filter pair is classified into one of eight filter distance groups (d1–d8).

Similarities in OI and CI differ among filter-distance groups in the model instance shown in Figure 2. In this model instance, some conv1 filters that did not develop clear structures are excluded from the analyses. From the 143 remaining filters, 10,153 filter pairs were constructed. The absolute difference in OI (*Δ*OI) between two filters differs among the eight filter-distance groups (*p* = 2.61 × 10^-61^, *n* = 10,153, Kruskal– Wallis test; Fig. 2C, left), and the *Δ*OI values of d1 pairs are smaller than those of the d2–d8 pairs (*p* = 1.80 × 10^-33^–2.20 × 10^-4^; Mann–Whitney U test). The absolute difference in CI (*Δ*CI) between two filters also differs among the filter-distance groups (*p* = 5.21 × 10^-57^; Fig. 2C, right), and the *Δ*CI values of d1 pairs are smaller than those of the d2–d8 pairs (*p* = 2.16 × 10^-29^–3.30 × 10^-5^).

Another salient feature of Figure 2A involves the similarities in preferred orientation (pO)^28^ among adjacent filters in the filter matrix. This tendency is visualized by plotting the pO values on the filter matrix (pO map; Fig. 2D, left). For example, filters around filter 155 prefer a vertical orientation, whereas filters around filter 219 prefer a horizontal orientation. The absolute difference in pO between two filters (*Δ*pO) represents the similarity in pO and takes values between 0° (the same orientation) and 90° (an orthogonal orientation). The analysis of the pO was limited to orientation-selective filters (*n* = 53, OI ≥ 0.285), which comprised 1,378 filter pairs. The absolute difference in pO (*Δ*pO) differs among filter-distance groups (*p* = 1.04 × 10^-9^, *n* = 1,378; Fig. 2E, left), and that of the d1 pairs is smaller than those of the d2–d8 pairs (*p* = 1.37 × 10^-10^–1.24 × 10^-4^).

Clustering of the preferred hue (pH)^28^ in the filter matrix is also observed (Fig. 2D, right). The absolute difference in pH between two filters (*Δ*pH) represents the similarity in pH and takes a value between 0° (the same color) and 180° (the opposite color). The analysis of pH was limited to color-selective filters (*n* = 53, CI ≥ 0.25), which comprised 1,378 filter pairs. (Note that the equality in color-selective and orientation-selective filter count (53) arises by chance and has no methodological significance.) The absolute difference in pH (*Δ*pH) differs among filter-distance groups (*p* = 0.0012, *n* = 1378; Fig. 2E, right), and that of the d1 pairs is smaller than those of the d7 and d8 pairs (*p* = 0.026 and 0.041, respectively).

The similarities in filter properties were further quantified using a variety of measures. The absolute difference in preferred spatial frequency (pSF)^28^ between two filters (*Δ*pSF) represents the similarity in pSF and takes a value between 0 (the same pSF) and 16 (the maximum pSF difference). The absolute difference in preferred spatial phase (pP)^28^ between two filters (*Δ*pP) represents the similarity in pP and takes values between 0° (in-phase) and 180° (anti-phase). The analysis of pP was limited to orientation-selective filters. The *Δ*pSF differs among filter-distance groups of the model instance in Figure 2 (*p* = 9.08 × 10^-34^, *n* = 10,153; Fig. 3, top row, left), and that of the d1 pairs is smaller than those of the d2–d8 pairs (*p* = 1.16 × 10^-25^–1.18 × 10^-5^). However, *Δ*pPdoes not differ among filter-distance groups in the model instance (*p* = 0.86, *n* = 1,378; Fig. 3, top row, right).

**Figure 3.**
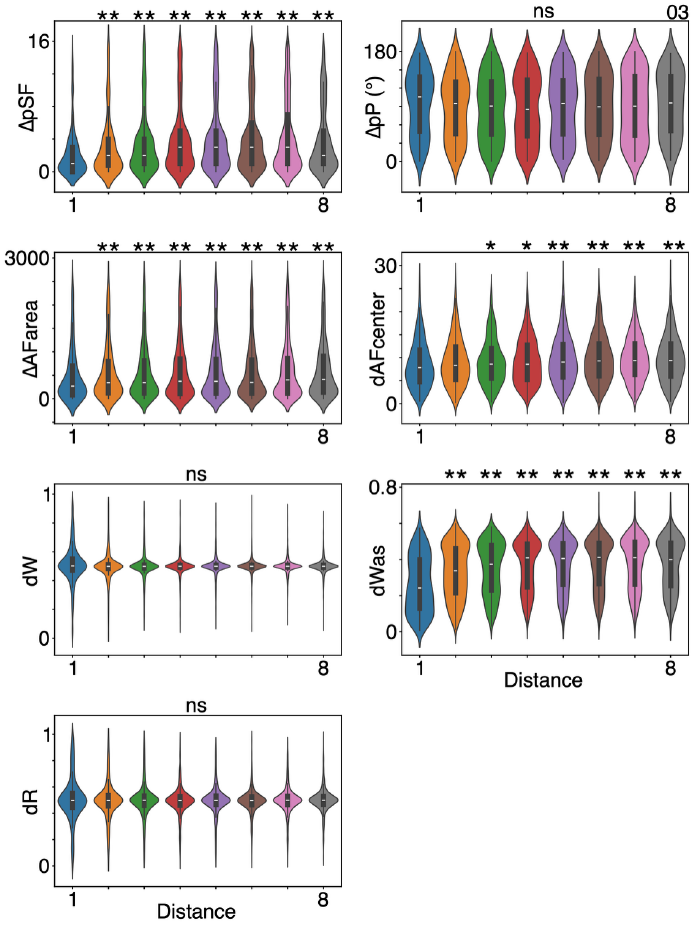
Comparisons of similarities in filter properties in the first convolutional layer (conv1) of tmcAlexNet across filter-distance groups for the representative model instance in Fig. 2. Comparisons of the absolute differences in the preferred spatial frequency (*Δ*pSF; top row, left), and preferred spatial phase (*Δ*pP; top row, right), and the area of the activation field (*Δ*AFarea; second row, left). Comparisons of the distances between the center position of the AFs (dAFcenter; second row, right). Comparisons of the distances of the pixel-by-pixel filter weights (dW; third row, right), amplitude spectrum of filter weights (dWas; third row, left), and set of responses (dR; bottom row, left). “ns” indicates non-significant differences across filter-distance groups. Other conventions are as in Fig. 2.

In this study, the conv1 filter was enlarged from the conventional size of 11 × 11 pixels to 33 × 33 pixels. In filters with conventional sizes, the region where a filter receives inputs, called the activation field (AF), extends over the entire 11 × 11 pixels region. However, when the filter size is increased to 33 × 33 pixels, the AFs of some of the conv1 filters of tmcAlexNet are not fully extended over the entire 33 × 33 pixels region. As a result, the area and center positions of AFs differ among filters (Supplementary Figure 4). For example, both filters 52 (AF area = 71.9 pixels^2^) and 217 (AF area = 177.5 pixels^2^) of the model instance in Figure 2 receive input from relatively small area, whereas filter 128 receives input from large area (AF area = 1171.8 pixels^2^). Furthermore, position of the AF center is not always located at the filter’s center, that is, (x, y) = (0, 0), but differs among filters. For example, filter 52 receives input from left-upper region (–5.18, 7.95), whereas filter 217 receives input from right-lower region (2.99, –7.83). This variability in AF structure was used to examine whether adjacent filters in the filter matrix have similar AF areas and center positions.

The similarity in AF area (*Δ*AFarea) differs among the filter-distance groups of the model instance shown in Figure 2 (*p* = 0.0059, *n* = 8,256; Fig. 3, second row, left). The *Δ*AFarea of the d1 pairs is smaller than those of the d2–d8 pairs (*p* = 2.71 × 10^-5^–0.0092). The distance between AF center position (dAFcenter) also differs among the filter-distance groups (*p* = 2.45 × 10^-7^, *n* = 8,256; Fig. 3, second row, right). The dAFcenter of the d1-pairs is smaller than those of the d3–d8 pairs (*p* = 2.17 × 10^-6^–0.036). The results suggest that filters with similar AF areas and center positions are clustered in the filter matrix of tmcAlexNet.

The filter-weight structure was quantified using two measures, and the distance for each measure was calculated to quantify the similarities between two conv1 filters in tmcAlexNet. One distance measure quantifies the pixel-by-pixel similarity in filter weights (dW), and the other quantifies the similarity in the amplitude spectrum calculated after a 2D discrete Fourier transform of the filter weights (dWas)^28^. The dW and dWas metrics take values between zero and one. If the filter structures are similar to each other, dW and dWas will be close to zero. The dW does not differ among filter-distance groups of the model instance shown in Figure 2 (*p* = 0.086, *n* = 10153; Fig. 3, third row, left). However, the dWas differs among filter-distance groups (*p* = 1.58 × 10^-50^, *n* = 10153; Fig. 3, third row, right), and that of the d1 pairs is smaller than those of the d2–d8 pairs (*p* = 7.33 × 10^-40^–2.71 × 10^-12^).

To further quantify the filter property, a set of filter responses to 1,000 images from the ImageNet database^31^ was obtained, and the distance measure (dR) between two filters in the set of responses was obtained^28^. The dR takes values between zero and one. If the filter responses are similar to each other, the dR is close to zero. The dR does not differ among the filter-distance groups of the model instance shown in Figure 2 (*p* = 0.34, *n* = 10,153; Fig. 3, bottom row, left).

Overall, these results suggest that filters with smaller filter distances tend to display similarity in some filter properties, and this similarity decreases with respect to filter distance in the conv1 filter matrix of tmcAlexNet.

Population analyses using all 16 model instances, each trained with random initial values, confirmed the relationships between filter distance and filter-property similarity observed in the representative model instance. In all the model instances, adjacent filters in the filter matrix tend to have similar properties (Supplementary Figure 5). The median absolute difference or distance for each filter metric across filter pairs was calculated for each filter-distance group of each model instance (Fig. 4). For *Δ*pSF, the mean absolute difference was calculated for each filter-distance group of each model instance because the median values did not capture the difference among filter distance groups. The median *Δ*OI, *Δ*CI, *Δ*pO, *Δ*pH, and mean *Δ*pSF increase with filter distance. The median dAFarea, dAFcenter, and dWas also increase with filter distance. However, the median *Δ*pP, dW, and dR do not increase with filter distance.

**Figure 4.**
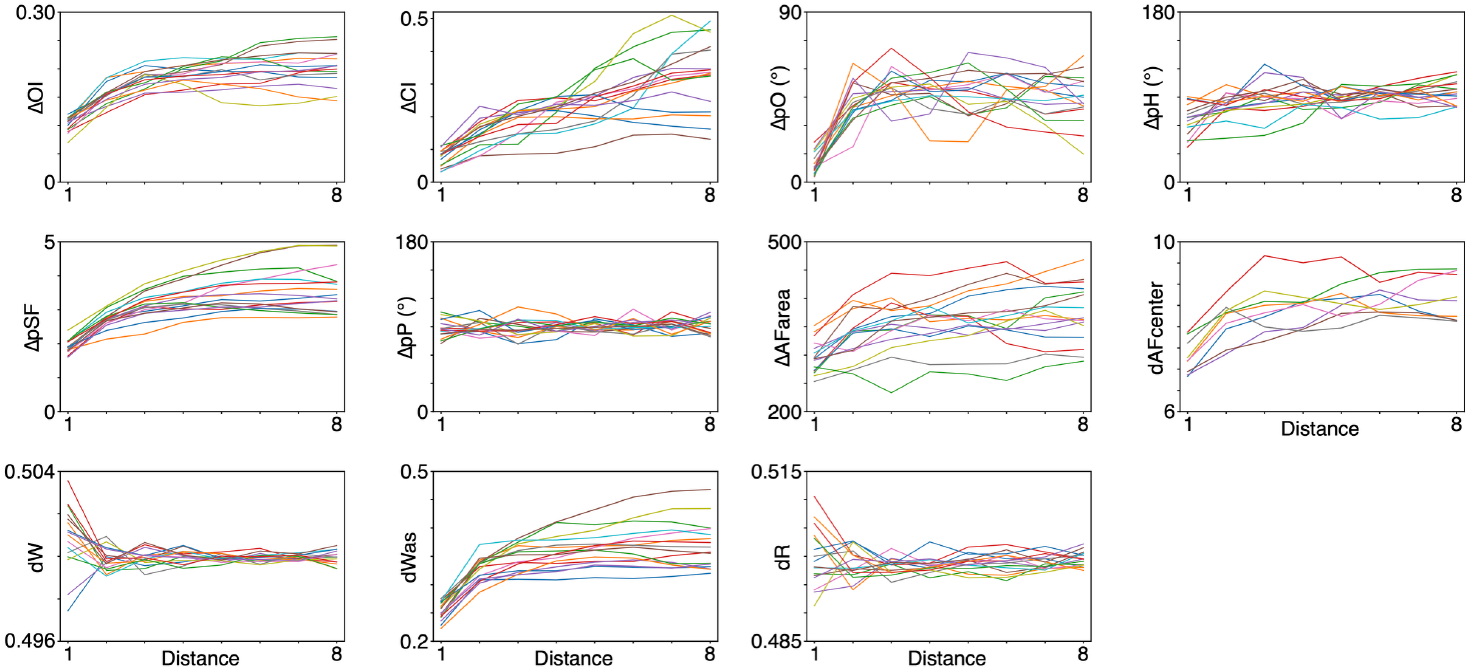
Relationships between the filter distance and filter-property similarity of the first convolutional layer (conv1) of tmcAlexNet for 16 model instances. Relationships between filter distance and the median absolute differences for the orientation index (ΔOI; top row, left), color index (ΔCI; top row, center left), preferred orientation (ΔpO; top row, center right), preferred hue (ΔpH; top row, right), preferred spatial frequency (ΔpSF; center row, left), and preferred spatial phase (ΔpP; center row, center left) across filter-distance groups. For ΔpSF, the mean absolute difference is plotted. Relationships between filter distance and the median absolute differences in the activation field area (ΔAFarea; center row, center right) and the median distance between activation-field center position (dAFcenter; center row, right). Relationships between the filter distance and median distance of pixel-by-pixel filter weights (dW; bottom row, left), amplitude spectrum of the filter weights (dWas; bottom row, center left), and set of responses (dR; bottom row, center right). Each line represents a model instance.

The relationship between the similarities in filter properties and filter distance was further quantified by calculating the correlation coefficients between them (Supplementary Figure 6). Positive correlations between filter distance and *Δ*OI, *Δ*CI, *Δ*pSF, and dWas (*p* < 0.05) were observed in 93.8% (15/16) instances. Distributions of the correlation coefficients between filter distance and *Δ*OI, *Δ*CI, *Δ*pSF, and dWas were shifted from zero in the positive direction (*p* = 3.05 × 10^-5^–9.16 × 10^-5^, Wilcoxon signed-rank test; Fig. 5), meaning that if the distance between two filters is small, they have similarities in these properties.

**Figure 5.**
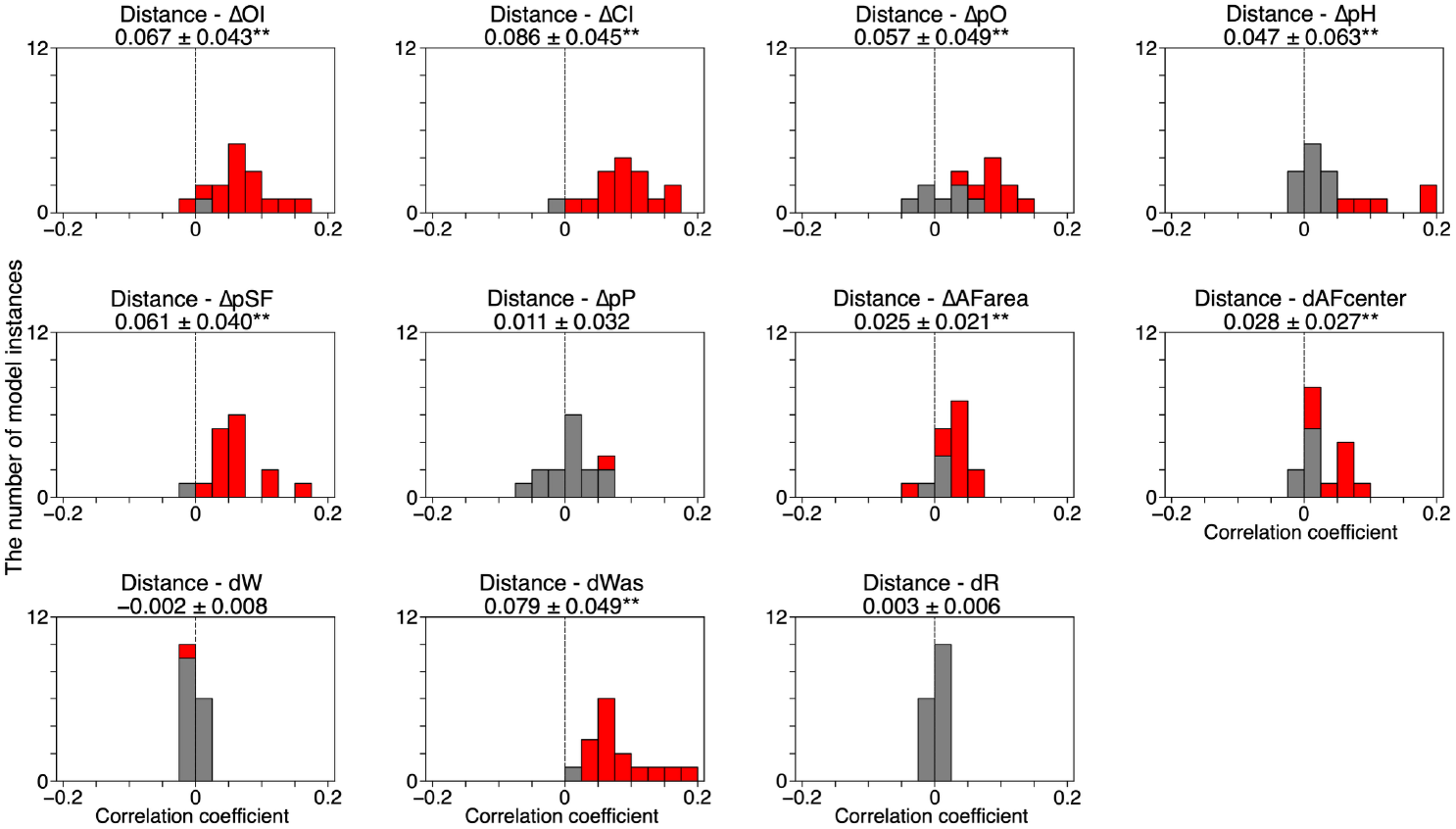
Frequency distributions of the correlation coefficients between filter distance and filter-property similarities of paired filters in the first convolutional layer (conv1) of tmcAlexNet for the 16 model instances. Correlation coefficients between filter distance and the absolute difference in orientation index (ΔOI; top row, left), color index (ΔCI; top row, center left), preferred orientation (ΔpO; top row, center right), preferred hue (ΔpH; top row, right), preferred spatial frequency (ΔpSF; center row, left), and preferred spatial phase (ΔpP; center row, center left). Frequency distributions of correlation coefficients between filter distance and the absolute difference in activation field area (ΔAFarea; center row, center right) and between filter distance and the distance in activation-field center position (dAFcenter; center row, right). Frequency distributions of the correlation coefficients between filter distance and the distance in pixel-by-pixel filter weights (dW; bottom row, left), amplitude spectrum of filter weights (dWas; bottom row, center left), and set of responses (dR; bottom row, center right). The red and gray columns indicate significant (*p* < 0.05) and non-significant correlation, respectively. Mean ± standard deviation of correlation coefficient across 16 model instances is provided in each panel. Double asterisks indicate a significant shift of the distribution from zero with *p* < 0.01.

Positive correlation was also observed between filter distance and *Δ*pO, *Δ*pH, *Δ*AFarea, and *Δ*AFcenter, and the distributions of these correlation coefficients were also shifted from zero in the positive direction (*p* = 3.05 × 10^-4^–0.0063, Wilcoxon signed-rank test; Fig. 5). Thus, similarity in these filter properties is related to filter distance, suggesting that these filter properties are organized in a topographic manner in the filter matrix of tmcAlexNet. However, the distributions of the correlation coefficients between filter distance and *Δ*pP, dW, or dR were not shifted from zero (*p* = 0.093-0.404; Fig. 5), suggesting that similarity in these filter properties is not related to filter distance.

### Extent of the functional module in conv1 of tmcAlexNet

The extent of the functional module was estimated from the OI and CI maps (see Fig. 2B) and the pO and pH maps of the conv1 filters of tmcAlexNet by calculating the 2D auto-correlogram (2DACG) of the maps (Fig. 6A). The 2DACG peaks of the OI and CI maps are broad, whereas those of the pO and pH are sharp. The full width at half height of the 2DACG peak was used as measure of the width of the 2DACG peak. In the representative case, the peak width of the 2DACG of the OI and CI maps are 3, and those of the pO and pH maps are 1.

**Figure 6.**
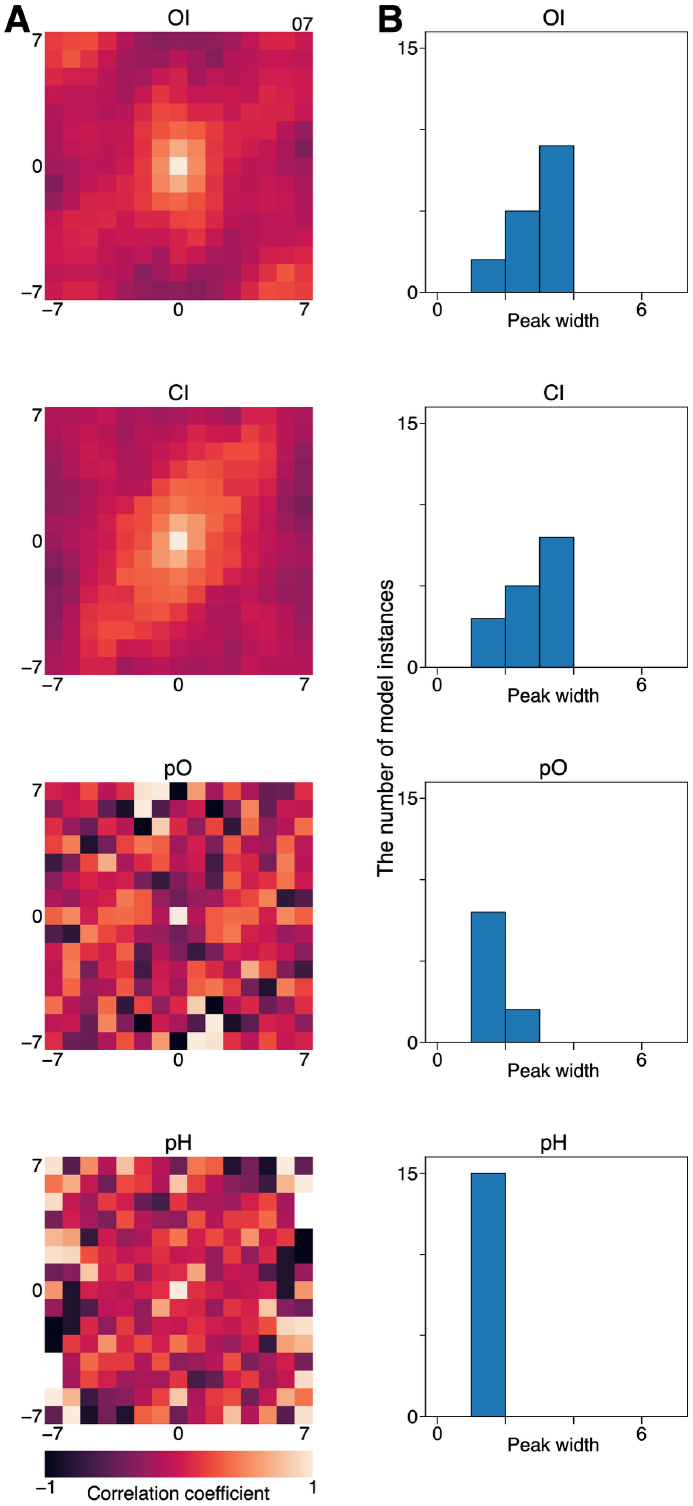
Comparisons of the peak widths of the two-dimensional auto-correlograms (2DACGs) of the orientation-index (OI), color-index (CI), preferred-orientation (pO), and preferred-hue (pH) maps of conv1 of tmcAlexNet. ***A***, 2DACGs for the OI (top), CI (second row), pO (third row), and pH (bottom) maps of a model instance. The correlation coefficients are color coded. ***B***, Frequency distributions of the peak widths of the 2DACGs for the OI (top), CI (second row), pO (third row), and pH (bottom) maps for the 16 model instances. The 2DACGs of the pO maps of six model instances and the pH map from one model instance were noisy and were excluded from the analysis.

The peak widths of the 2DACGs differed among the different types of functional maps (*p* = 2.85 × 10^-4^, 16 model instances, Friedman test for repeated samples). Those of the OI and CI maps are 2.4 ± 0.7 (mean ± standard deviation, *n* = 16) and 2.3 ± 0.77 (*n* = 16), respectively. Those of the pO and pH maps are 1.2 ± 0.4 (*n* = 10) and 1 ± 0 (*n* = 15), respectively. The fact that the peak width of the 2DACG of the index maps, i.e., OI and CI maps (2.38 ± 0.74), is larger than that of the preference maps, i.e., pO and pH maps (1.08 ± 0.27, *p* = 6.68 × 10^-5^), suggests that the extent of the preference module is smaller than that of the index module, as expected.

### Relationships between classification accuracies and similarities in filter properties

Because relationships exist between filter distance and the similarity of filter properties in conv1 of tmcAlexNet, such organization may play an important role in the classification of input images. If this is the case, the degree of positive correlation between filter distance and filter-property similarity is likely to be related to classification accuracy. To examine this possibility, we calculated correlations between the top-5 accuracy and correlation coefficients between filter distance and filter-property similarity using all 16 model instances. Here, analysis was limited to the eight filter-similarity measures that showed a positive relationship with filter distance, i.e., *Δ*OI, *Δ*CI, *Δ*pO, *Δ*pH, *Δ*pSF, dAFarea, dAFcenter, and dWas.

Top-5 accuracy is positively related to the correlation coefficient between filter distance and ΔCI (*r* = 0.54; Fig. 7, top row, center). Top-5 accuracy is also positively related to the correlation coefficient between filter distance and dWas (*r* = 0.54; Fig. 7, bottom row, center). The results indicate that if similarity in CI or Was decreases with respect to the filter distance in a model instance, that model instance achieved higher classification accuracy than model instances in which the relationships between similarity in CI or Was and filter distance was weak.

**Figure 7.**
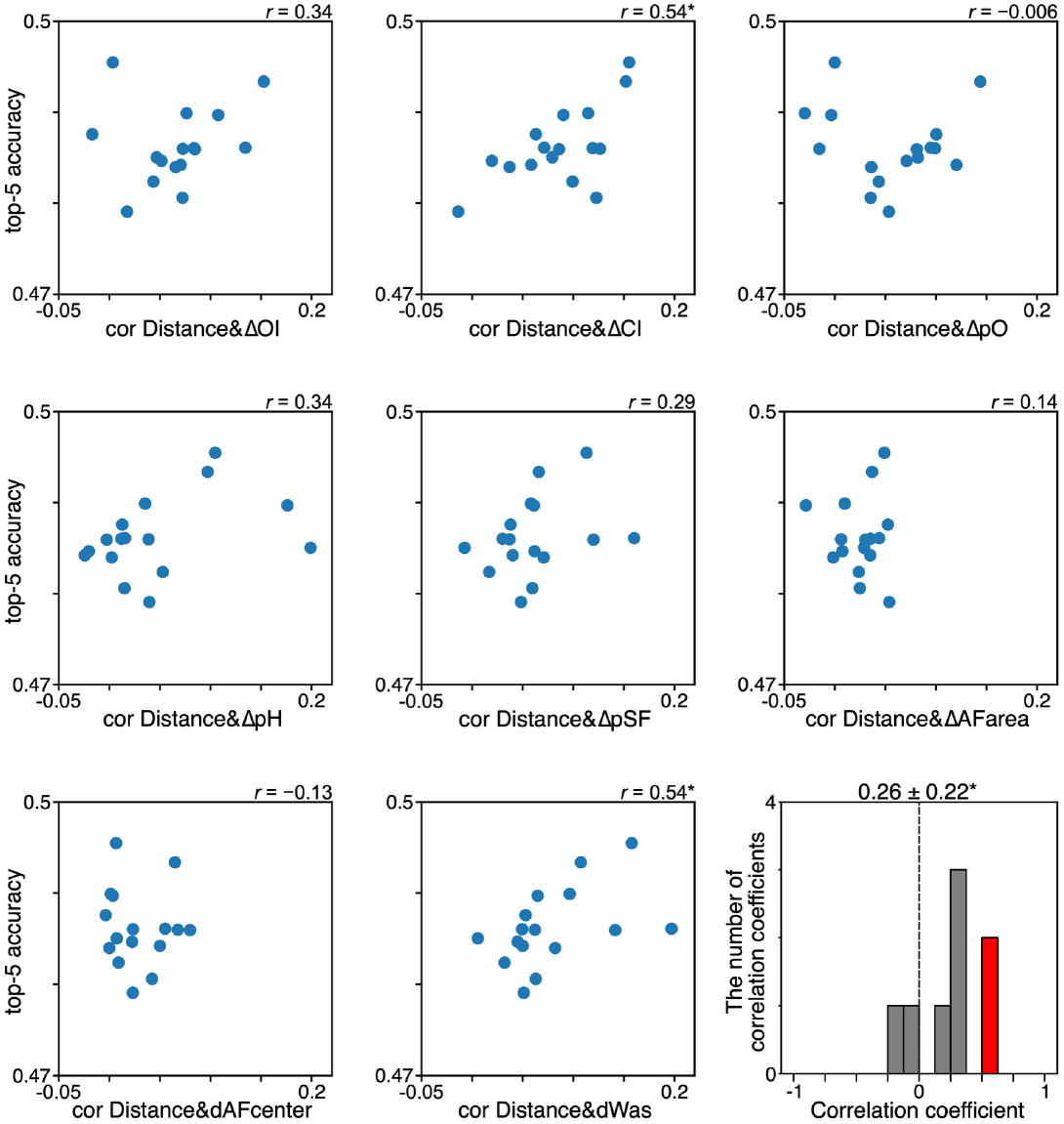
Relationships between top-5 accuracy and the correlation coefficient between filter distance and filter-property similarity of conv1 of tmcAlexNet. Relationships between top-5 accuracy and the correlation coefficient between filter distance and the absolute difference in orientation index (cor Distance&*Δ*OI; top row, left), absolute difference in color index (cor Distance&*Δ*CI; top row, center), absolute difference in preferred orientation (cor Distance&*Δ*pO; top row, right), absolute difference in preferred hue (cor Distance&*Δ*pH; center row, left), absolute difference in preferred spatial frequency (cor Distance&*Δ*pSF; center row, center), absolute difference in activation field area (cor Distance&*Δ*AFarea; center row, right), distance in activation-field center position (cor Distance&dAFcenter; bottom row, left), and distance using the amplitude spectrum of the filter weights (cor Distance&dWas; bottom row, center). Single points in each graph represent a model instance, and 16 points are plotted. The correlation coefficient (Spearman’s rank correlation) is provided for each panel. In addition, frequency distribution of the correlation coefficient is provided (bottom row, right). The red and gray columns indicate significant (*p* < 0.05) and non-significant correlation, respectively. Mean ± standard deviation of the correlation coefficients is provided. Single asterisks indicate significant shift of distribution from zero with *p* < 0.05.

Weak but clear positive relationships between top-5 accuracy and the correlation coefficients between filter distance and *Δ*OI, *Δ*pH, or *Δ*pSF were also observed (Fig. 7). Distributions of the correlation coefficients between the top-5 accuracy and the correlation coefficient between filter distance and filter-property similarity were shifted from zero in a positive direction (0.26 ± 0.22 (mean ± standard deviation), *n* = 16; *p* = 0.039, Wilcoxon signed-rank test; Fig. 7), suggesting that the relationships between filter distance and filter-property similarity are important for the classification of input images.

The relationship between top-5 accuracy and the filter-property similarity of the d1 pairs were also examined to clarify the importance of the degree of clustering on the classification of input images (Supplementary Figure 7). If similarity in the d1 pairs plays an important role in the classification of input images, the correlation between difference or distance measures and top-5 accuracy should be negative. However, negative significant correlation was not observed, and the distribution of correlation coefficients was not shifted from zero (–0.02 ± 0.23, *n* = 16; *p* = 1.0). Thus, degree of clustering is not related to top-5 accuracy.

### Integration of the outputs from orientation selective conv1 filters and color selective conv1 filters produces a variety of filter properties in conv2

Conv1 filters with similar properties are clustered in most of the filter matrix, and conv2 filters integrate the outputs from conv1 filters with similar properties (Supplementary Figure 8). However, because in some parts of the filter matrix, orientation-selective filters are adjacent to color-selective filters, some conv2 filters may integrate the outputs from both orientation selective conv1 filters and outputs from color selective conv1 filters. To examine whether and how conv2 filters integrate outputs from conv1 filters with different properties, stimulus images that maximally activates (MA) conv2 filters were visualized^27^.

Conv2 filters in a stream of a model instance receive outputs from two orientation-selective conv1 filters and two color-selective conv1 filters (Fig. 8). Although these conv2 filters can integrate information from different submodalities, they may receive information only from a single submodality by adjusting their input weights. Indeed, the MA images of some conv2 filters in the stream primarily consist of information from a single submodality; for example, the 4th filter (top row, right, predominantly orientation) and 12th filter (bottom row, right, predominantly color) in Figure 8B. However, the MA images of other conv2 filters consist of a combination of orientation-selective and color-selective components (for example, the 5th filter (center row, left)) suggesting integration of different submodality information by a single conv2 filter. Thus, conv2 filters acquire a variety of properties by integrating the outputs from orientation-selective conv1 filters and color-selective conv1 filters that are adjacent to each other in the filter matrix with a variety of weights.

**Figure 8.**
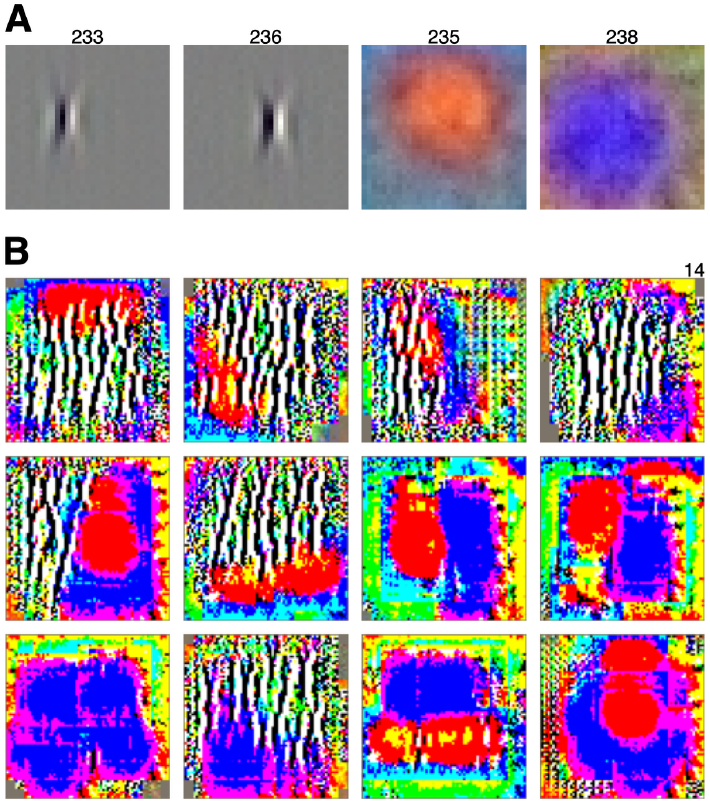
Examples of images that maximally activate (MA) filters in the second convolutional layer (conv2) of mcAlexNet. *A*. Visualization of filter weight for four filters (filters 233, 236, 235, and 238) in the first convolutional layer (conv1) connecting to a set of 12 conv2 filters in a stream. *B*. MA images for the 12 conv2 filters. MA images were calculated for filter-units having receptive field at the center.

## Discussion

In tmcAlexNet, the topographic organization of conv1 filters emerged spontaneously. Filter weights were initially set to random values and there were no constraints on the input weights for the conv1 filters. Therefore, the filters were able to adopt any filter structure irrespective of the location in the filter matrix. However, filter pairs with smaller filter distances tend to display similar properties and this similarity decreases with filter distance.

The spontaneous emergence of topographic organization in the filter matrix of tmcAlexNet is consistent with a self-organization model of a columnar structure in the primary visual cortex^32^, and the topographic organization of filter properties can emerge spontaneously without providing a detailed connection diagram.

The constrained connections from conv1 filters to conv2 filters are likely to contribute to the spontaneous emergence of topographic organization in the filter matrix. The filter matrix of conv1 filters is constructed based on the number of target-conv2 filters shared by the conv1 filters, and the connections from conv1 to conv2 changes gradually across the filter matrix. In other words, adjacent conv1 filters in the filter matrix connect to the same target filters in conv2. Because there are no constraints other than the connections between conv1 and conv2, the constrained connections induce the topographic organization of conv1 filters across the filter matrix.

The spontaneous emergence of topographic organization in the conv1 filter matrix of tmcAlexNet enables conv2 filters to integrate similar information from conv1 filters. Topographic organization is observed for OI and CI in the conv1 filter matrix of tmcAlexNet, meaning that conv2 filters integrate information about the same type of visual submodality. This type of integration restricts the analysis of input images by the conv2 filters to a certain visual submodality and improves resolution in the submodality by circumventing combinatorial explosion^28^.

Topographic organization was also observed for pO, pH, pSF, AF area, and AF center position, indicating that conv2 filters integrate similar orientation, color, SF, AF area, and AF center position information. Conv2 filters may compare outputs from conv1 filters encoding similar stimulus parameters to enhance discriminability in these parameters.

In contrast, topographic organization was not observed for pP. This observation suggests that integrations are performed using the outputs from filters with similar pO, pH, and pSF, but different pP. By integrating this type of information, representations for orientation, color, and SF that are invariant to spatial phase can be constructed.

The topographic organization of OI, CI, pO, pH, pSF, AF area, and AF center position observed in the filter matrix of conv1 in tmcAlexNet is similar to that observed in V1 of primates. Orientation-selective neurons are clustered in inter-blob regions and color-selective neurons are clustered in blob regions^4-10^. Furthermore, orientation-selective neurons are clustered according to the preferred orientation, which changes smoothly across the cortex^11-14^. In the V1 of primates, neurons preferring modulation in the red–green axis or the blue–yellow axis are clustered in local regions^34^. Studies using two-photon imaging revealed the clustering of neurons according to preferred color^16^. Preferred spatial frequency also changes smoothly across the cortex^17, 18^. RF position is smoothly mapped across the cortex, i.e., the retinotopic organization, and RF area is related to RF position^1-3^.

Failure to find topographic organization in pP in the filter matrix of conv1 in tmcAlexNet is also similar to observations of the cat primary visual cortex, in which neighboring neurons often prefer a variety of spatial phases^33^. To the best of the author’s knowledge, the topographic organization of preferred spatial phase has not been reported.

Topographic organizations in V1 enable two seemingly conflicting requirements. By integrating outputs from neighboring neurons that carry similar but not identical information, target neurons can enhance stimulus discriminability and acquire invariant response properties. At the same time, by integrating outputs from neighboring neurons at the border of functional modules each carrying different information, target neurons can acquire a filter property that is not observed in the source neurons. Similarly, some neurons in V2 of primates are selective to more than one visual submodality^35^.

Although the present study focused on conv1 filters, which have properties similar to those of V1 neurons, topographic organization is also observed in higher visual cortical areas, such as V4^36, 37^ and the inferior temporal cortex^38-41^ of primates. In tmcAlexNet, filters in the higher layers are not placed in a filter matrix and receive constrained inputs. Thus, topographic organizations in the higher layers are difficult to study using the current model architecture. In future studies, this issue should be addressed.

## Methods

tmcAlexNet consisted of five hierarchically organized convolutional layers (conv1–conv5) and three pooling layers. Conv1 contained 256 filters. Conv2 contained 256 streams (s1–s256), each equipped with 12 filters. Conv3 contained 64 streams (s1–s64), each equipped with 96 filters. Conv4 contained 16 streams (s1–s16), each equipped with 256 filters. Finally, conv5 contained four streams (s1–s4), each equipped with 1024 filters (Fig. 1).

The conv1 filter was expanded from the conventional area of 11 × 11 pixels to 33 × 33 pixels to reveal the relationship between filter distance and similarity in AF area and AF center position.

Each conv2 filter received converging inputs that came from four conv1 filters. The connections between conv1 and conv2 were designed so that each conv1 filter shared a target with eight other conv1 filters. The filters at the edges of the filter matrix wrap across; that is, the edge filters share a target with the filters at the opposite edge. As a result, even the filters at the edges share a target with eight other filters (Supplementary Figure 1). Each conv3 filter received converging inputs that came exclusively from 48 filters in four conv2 streams. Each conv4 filter received converging inputs that came exclusively from 384 filters in four conv3 streams. Finally, each conv5 filter received converging inputs that came exclusively from 1024 filters in four conv4 streams. The outputs from the four conv5 streams were concatenated and fed into FCs and then passed to the output layer for classification.

tmcAlexNet was trained to classify 1,000 object-image categories using the ImageNet database^31^ following the procedure in a previous report^27, 28^ except for the number of training epochs, which was 30 in this study. This is because the filter-weight structures of conv1 filters develop quickly and become almost stable after six epochs (Supplementary Figure 9). Training was performed 16 times, and each model instance was trained using randomly initialized parameters. tmcAlexNet was also trained for image representation using VICReg^42^, and results similar to those for the classification training were obtained (Supplementary Figure 10).

The orientation selectivity of filters in conv1 was quantified using OI^28^. The OI value was between zero and one, with larger values reflecting stronger orientation selectivity. *Δ*OI was calculated as the absolute difference in OI between two filters and represented similarity in OI. The color selectivity of each filter in conv1 was quantified using CI^27^, which was between zero and one, with larger values reflecting greater color selectivity. *Δ*CI was calculated as the absolute difference in CI between two filters and represented similarity in CI.

The preferred orientation, spatial frequency, spatial phase, and color of each filter in conv1 were quantified using pO pSF, pP, and pH^28^. Vertical and horizontal orientation were 0° and 90°, respectively. *Δ*pO was calculated as the absolute difference in pO between two filters and represented similarity in pO. *Δ*pO took values between 0° (the same orientation) and 90° (the orthogonal orientation). *Δ*pSF was calculated as the absolute difference in pSF between two filters and represented similarity in pSF. *Δ*pSF took values between 0 (the same pSF) and 16 (the maximum pSF difference). *Δ*pP was calculated as the absolute difference in pP between two filters and represented the similarity in pP. *Δ*pP took values between 0° (in-phase) and 180° (anti-phase). Color preference was quantified using the hue value in the HSV color space as pH. *Δ*pH was calculated as the absolute difference in pH between two filters and represented the similarity in pH. *Δ*pH took values between 0° (same color) and 180° (opposite color). The analysis of pO and pP was limited to orientation-selective filters and that of pH was limited to color-selective filters. If a filter did not develop any structure, as determined by the variance in the weight values, the filter was excluded from the analyses^27^.

To quantify the area and center position of the AF, a 2D Gaussian kernel was fitted to the filter-weight structure. Before fitting, the absolute values of the filter weights were calculated. If the fitting diverged, the filter was excluded from the analysis. The area within two standard deviations of the Gaussian kernel was defined as the AF area and the center position of the Gaussian kernel was defined as the AF center position (Supplementary Figure 4).

The filter-weight structure was evaluated using two measures^28^, W (the pixel-by-pixel filter weight) and Was (the amplitude spectrum calculated after a 2D discrete Fourier transform of the filter weight). From these measures, dW and dWas were calculated using the Pearson’s *r* values of W and Was for two filters, respectively. dW and dWas took values between zero and one. Similar filter structures yielded dW and dWas values close to zero.

The filter responses were examined using 1,000 images^28^. One image was randomly selected from each image category of the ImageNet database such that 1,000 images were obtained from the 1,000 categories ^31^. The similarity between sets of responses of two filters was evaluated by calculating a distance measure (dR) using the Pearson’s *r* value of the set of responses between the filters. dR took values between zero and one. Similar responses to the stimulus set gave a dR value close to zero. dR was examined using filter-units that had a receptive field at the center.

The 2DACG of OI and CI maps and those of pO and pH maps were calculated to estimate the extent of the functional module. For OI and CI maps, the 2DACG was obtained by calculating Pearsons’s correlation coefficient. For pO and pH maps, the 2DACG was obtained by calculating the circular correlation coefficient (Jammalamadaka–Sengupta correlation coefficient)^43^. To apply circular statistics, pO, which takes values between 0°-180°, was doubled (0°-360°) before 2DACG calculation. The full width at half height of the 2DACG peak was used to represent the module extent and was calculated by separately calculating the full width at half height of the 2DACG peak in the x and y directions and then taking the average of the two peak-width values. If the 2DACG peak height was smaller than the mean of the 2DACG + two × the standard deviation of the 2DACG, indicating that the 2DACG was noisy, it was excluded from the analysis.

### Statistical analysis

All data were pooled for statistical analyses. Analyses were performed using Python libraries (matplotlib, numpy, pandas, pingouin, scikit-learn, scipy, and seaborn). The statistical tests used in this study were the Mann–Whitney U test (two-tailed), Wilcoxon signed-rank test (two-tailed), and the Friedman test for repeated samples. All *r* values were Spearman’s rank correlation unless otherwise stated. The statistical threshold for *p-* values was set at 0.05. Median values were calculated to represent a population except for the SF and ACG peaks, in which the median did not capture the difference between groups and the mean value was calculated. For population measures of *r* values, the mean ± standard deviation was calculated.

## Data availability

Parts of the datasets generated and/or analyzed during the current study are available at the Osaka University Knowledge Archive (https://doi.org/10.60574/103843). The remaining data are available from the corresponding author on reasonable request.

## Code availability

Parts of the computer code used during the current study are available at the Osaka University Knowledge Archive (https://doi.org/10.60574/103843). The remaining computer code is available from the corresponding author on reasonable request.

## Acknowledgments

This work was supported by JSPS KAKENHI Grant Number 25K09847. We thank Kimberly Moravec, PhD, from Edanz (https://jp.edanz.com/ac) for editing a draft of this manuscript.

## Additional information

### Author contributions

HT designed the research, conducted the experiments, analyzed the data, and wrote the paper.

