## Supplemental Figures for "Spontaneous emergence of topographic organization in a multistream convolutional neural network"

20260224

**Hiroshi Tamura**[1, 2*]

[1]Graduate School of Frontier Biosciences, The University of Osaka, Suita, Osaka 565-0871, Japan

[2]Center for Information and Neural Networks, Suita, Osaka 565-0871, Japan

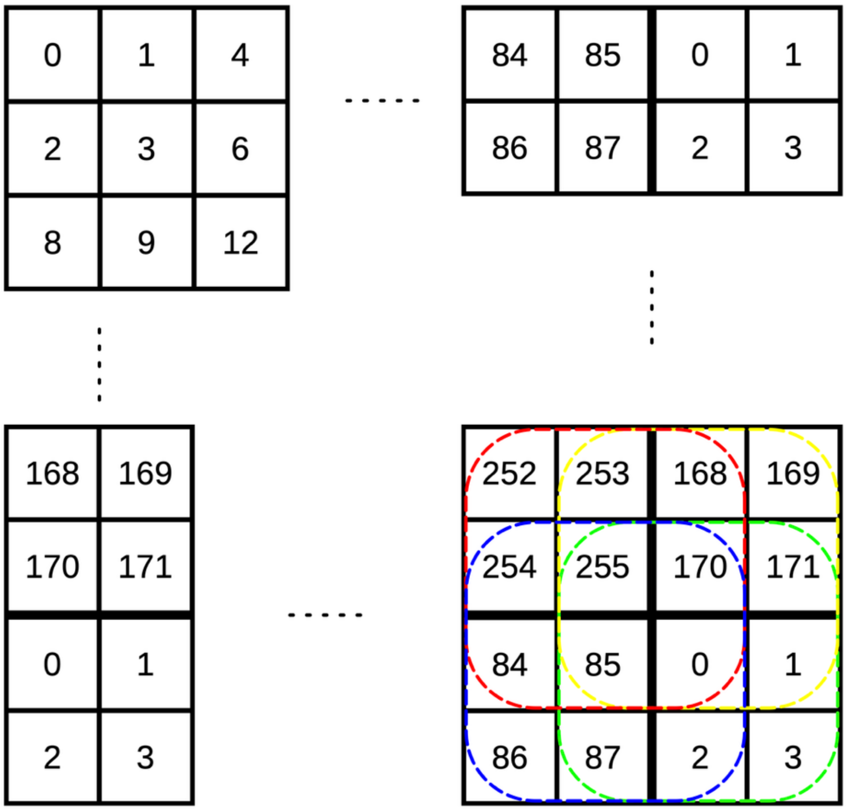


**Supplementary Figure 1.** Output connection patterns of the first convolutional layer (conv1) in a multistream convolutional neural network (tmcAlexNet), which was designed such that a filter shares targets with eight other filters. For example, filter 3 shares a conv2 target with filters 0–2, 4, 6, 8, 9, and 12. These eight filters were placed around filter 3. In this way, all 256 conv1 filters were placed in the filter matrix. Filters at the edge of the filter matrix share a target with the filters at the other edge. Hence, each filter shares a target with eight other filters. For example, filter 0 shares target with filters 1–3, 85, 87, 170, 171, and 255 (green-dashed border); filter 85 shares a target with filters 0, 2, 84, 86, 87, 170, 254, and 255 (blue-dashed border); filter 170 shares a target with filters 0, 1, 85, 168, 169, 171, 253, and 255 (yellow-dashed border); and filter 255 shares a target with filters 0, 84, 85, 168, 170, and 252–254 (red-dashed border).

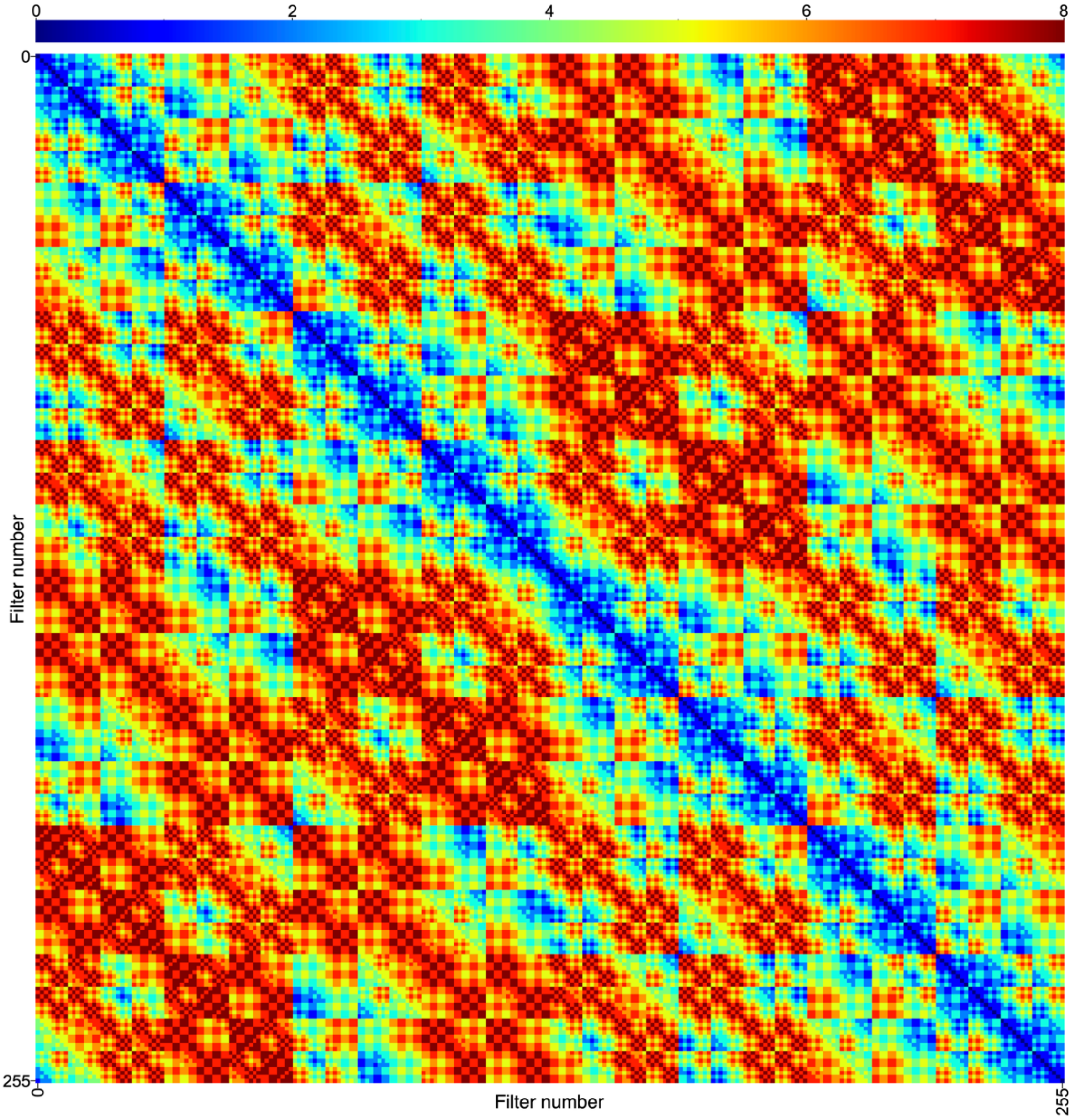


**Supplementary Figure 2.** Color-coded matrix of the filter distances between pairs of filters in conv1 of tmcAlexNet. The filter distance is color coded.

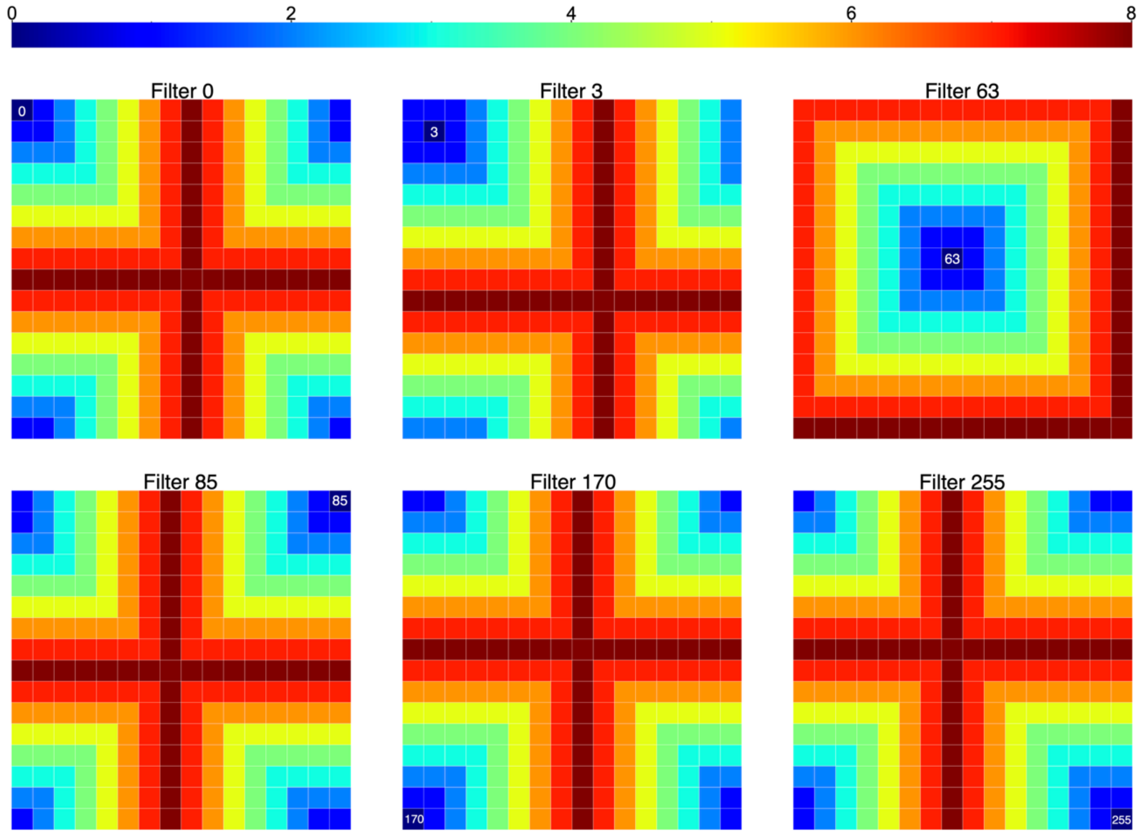


**Supplementary Figure 3.** Six examples of the color-coded matrix of the filter distances between filters, which are indicated by white numbers in the matrix, and other 255 filters in conv1 of tmcAlexNet. The filter distance is color coded.

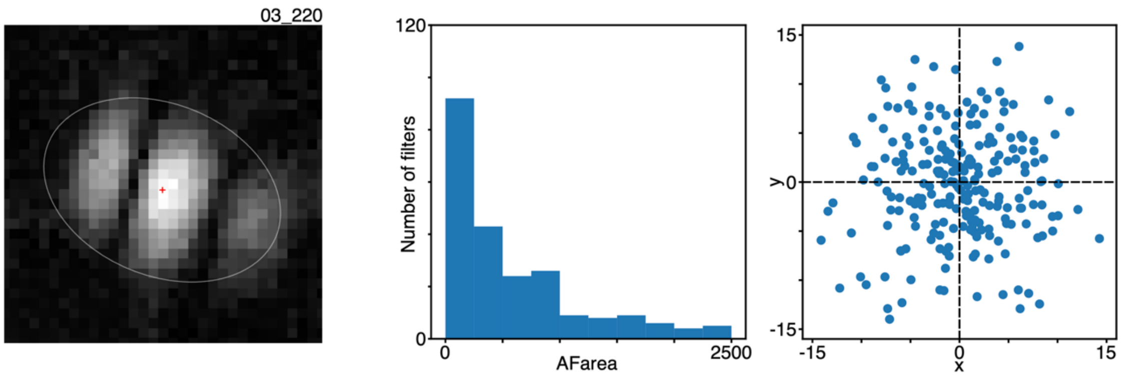


**Supplementary Figure 4.** Quantification of the activation field (AF) area and AF center position of filters in conv1 of tmcAlexNet. A two-dimensional Gaussian kernel was fitted to the filter-weight structure (left). Before fitting, the absolute values of the filter weights were calculated. The area within two standard deviations of the Gaussian kernel was defined as the AF area (white ellipse) and the center position of the Gaussian kernel (red cross) was defined as the AF center position. Frequency distribution of the AF areas (center) and spatial distribution of the AF center positions (right) of the conv1 filters of a model instance.

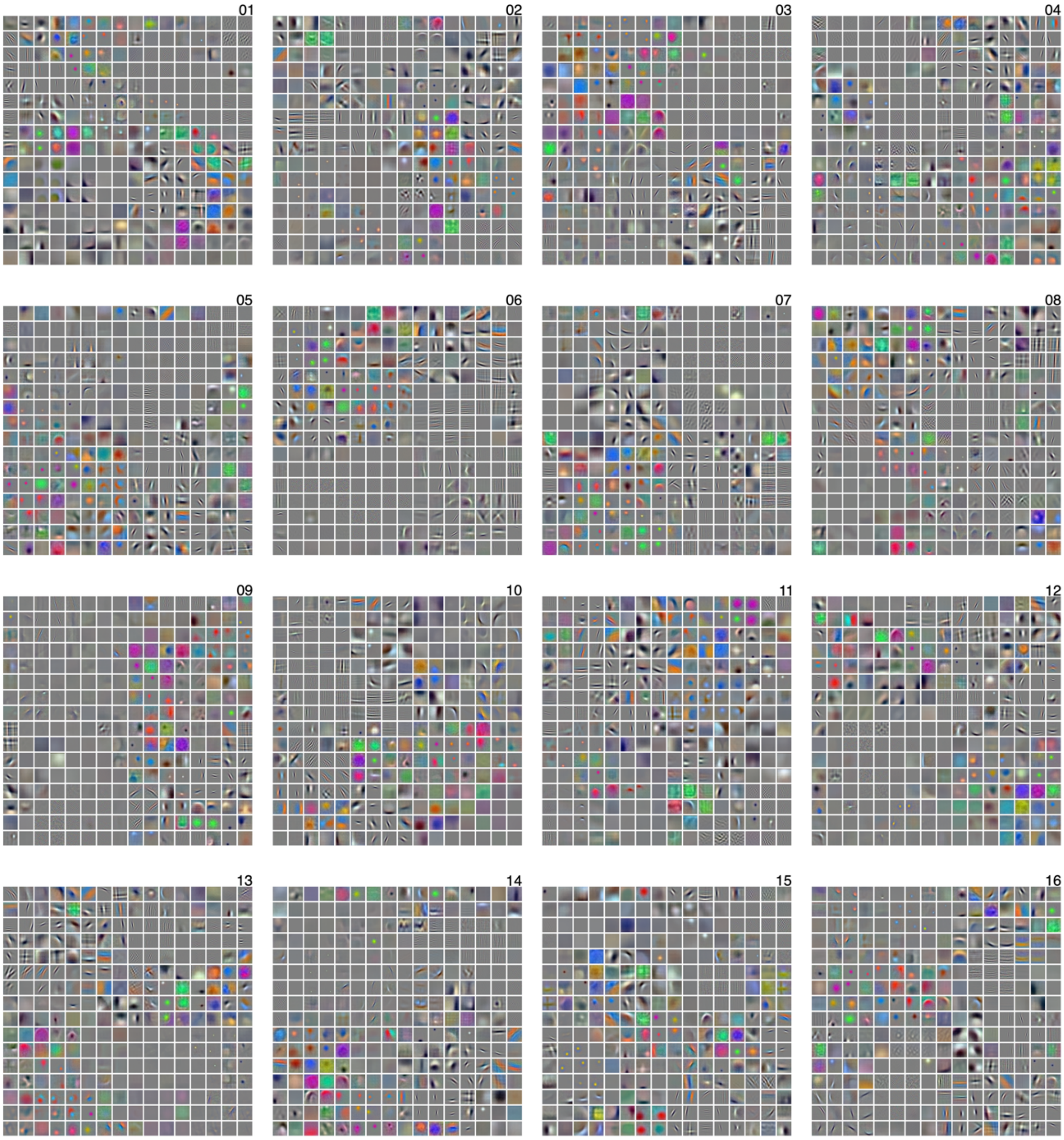


**Supplementary Figure 5.** Spatial arrangement of filters in conv1 of 16 model instances of tmcAlexNet. The weights for the conv1 filters are visualized. Filters are placed according to the filter matrices (see Figure 2A, right). The minimum and maximum weight values are scaled between 0 and 255 for visualization.

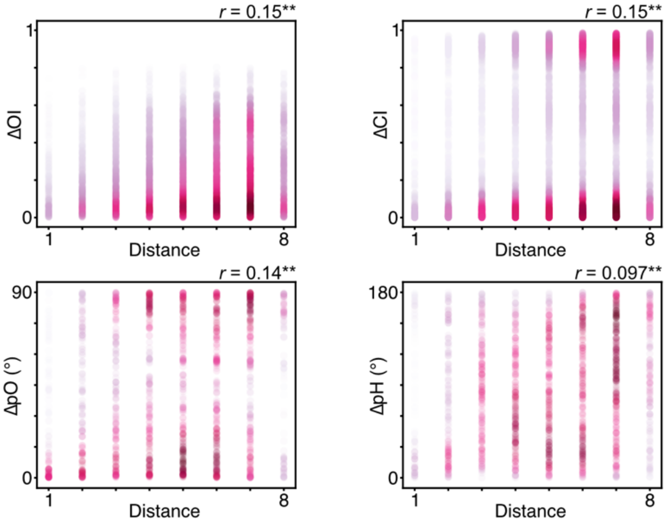


**Supplementary Figure 6.** Examples of the correlation between the filter distance and filter-property similarities of the paired filters in conv1 of tmcAlexNet. Relationships between filter distance and absolute differences in the orientation index (ΔOI; top left), color index (ΔCI; top right), preferred orientation (ΔpO; bottom left), and preferred hue (ΔpH; bottom right). Single points in each graph represent a paired filter; 10,153 points are plotted for ΔOI and ΔCI; 1,378 points are plotted for ΔpO; and 1,378 points are plotted for ΔpH. The correlation coefficient (Spearman's rank correlation) is provided for each panel. Double asterisks indicate significant correlation with $p < 0.01$.

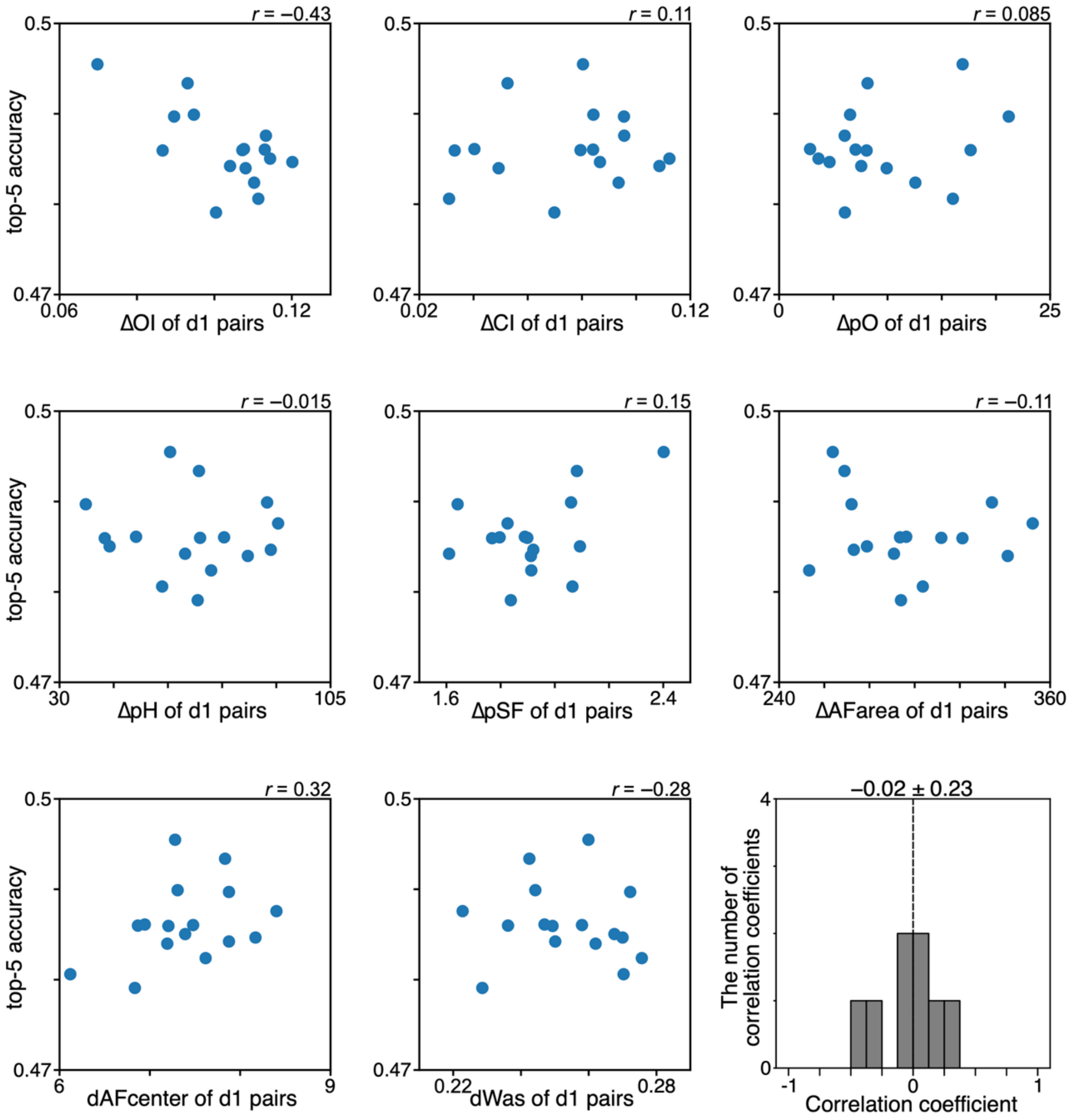


**Supplementary Figure 7.** Relationships between the top-5 accuracy and filter-property similarity of the adjacent filters (d1 pairs) in conv1 of tmcAlexNet. Relationships between the top-5 accuracy and the median absolute difference in orientation index (*Δ*OI of the d1 pairs; top row, left), median absolute difference in color index (*Δ*CI of the d1 pairs; top row, center), median absolute difference in preferred orientation (*Δ*pO of the d1 pairs; top row ,right), median absolute difference in preferred hue (*Δ*pH of the d1 pairs; center row, left), mean absolute difference in preferred spatial frequency (*Δ*pSF of the d1 pairs; center row, center), median absolute difference in activation field area (*Δ*AFarea of the d1 pairs; center row, right), median distance in activation-field center position (dAFcenter of the d1 pairs; bottom row, left), and median distance using the amplitude spectrum of the filter weights (dWas of the d1 pairs; bottom row, center). Single points in each graph represent a model instance; 16 points are plotted. The correlation coefficient (Spearman's rank correlation) is provided for each panel. In addition, the frequency distribution of the correlation coefficients is provided (bottom row, right). The gray columns indicate non-significant correlation ($p \geq 0.05$). The mean ± standard deviation of the correlation coefficients is provided. No asterisk indicates a non-significant ($p \geq 0.05$) shift in distribution from zero. There was a negative correlation between the median ΔOI and top-5 accuracy ($r = -0.43$, top left), which indicates importance of similarity in the d1 pairs in the classification of input images, but this correlation was not significant. Negative significant correlation was not observed in any similarity measures, and the distribution of the correlation coefficient was not shifted from zero.

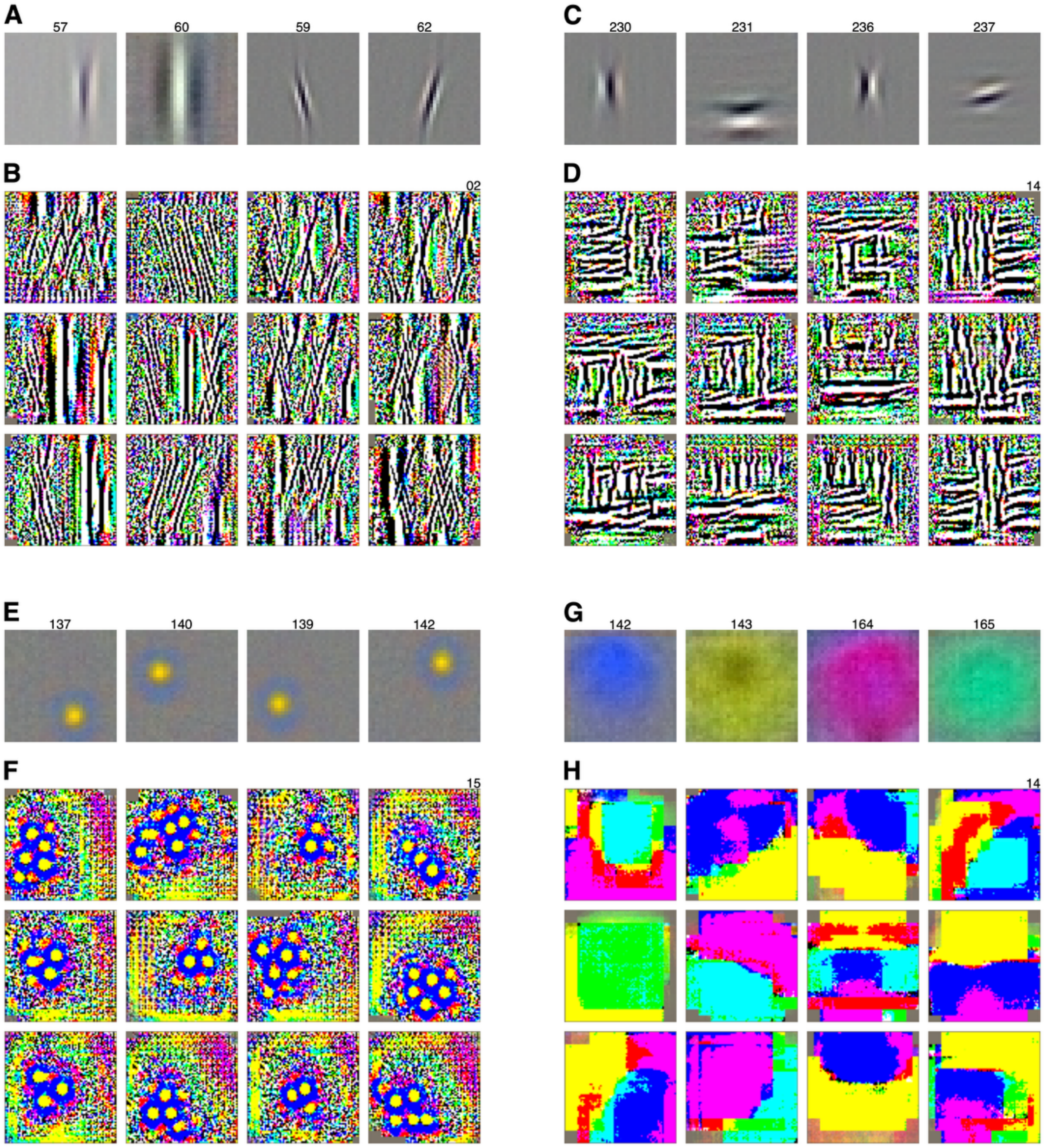


**Supplementary Figure 8.** Examples of images that maximally activate (MA) the filters in the second convolutional layer (conv2) of tmcAlexNet. *A*. Visualization of the filter weights for four orientation-selective filters in conv1 with similar preferred orientations. *B*. MA images for the 12 conv2 filters in a stream that integrate the outputs from the four filters in A. *C*. Visualization of the filter weight for four orientation-selective conv1 filters with different preferred orientations. *D*. MA images for the 12 conv2 filters in a stream that integrate the outputs from the four filters in C. *E*. Visualization of the filter weights for four color-selective conv1 filters with similar preferred colors. *F.* MA images for the 12 conv2 filters in a stream that integrate the outputs from the four filters in E. *G*. Visualization of the filter weights for four color-selective conv1 filters with different preferred colors. *H.* MA images for the 12 conv2 filters in a stream that integrate the outputs from the four filters in G. The MA images were calculated for filter units that had the receptive field at the center.

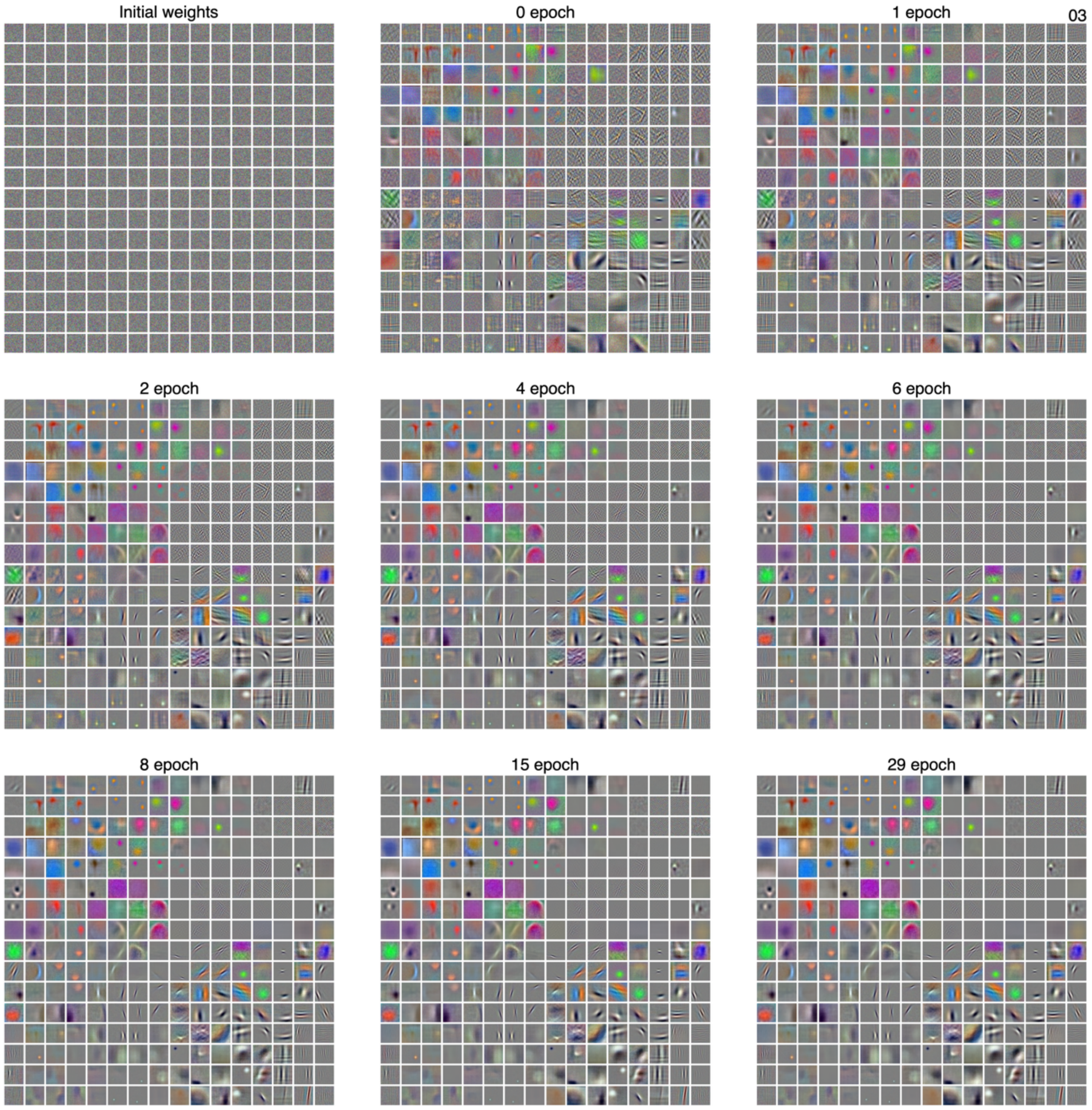


**Supplementary Figure 9.** Examples of developmental changes in the filter weights of conv1 for a representative model instance of tmcAlexNet. Note that the filter structures of epoch 0 (immediately after the first training epoch) already look similar to those of the last epoch (epoch 29) and those of epoch 6 are almost identical to those of epoch 29.

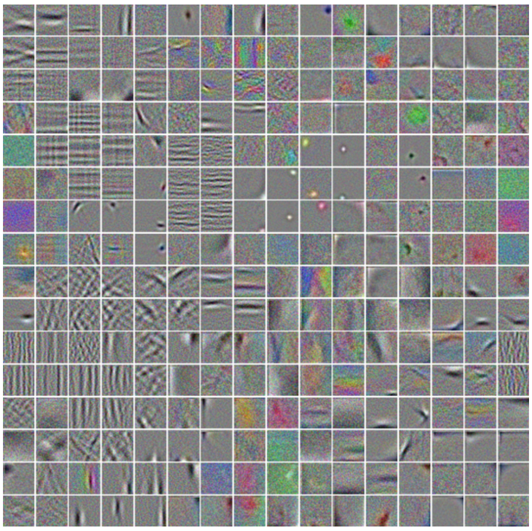

**Supplementary Figure 10.** Spatial arrangement of the filters in conv1 of a tmcAlexNet trained for image representation using VICReg for 200 epochs. tmcAlexNet was used as the backbone. The weights for the conv1 filters are visualized, and filters were placed according to the filter matrix (see Figure 2A, right). The minimum and maximum weight values are scaled between 0 and 255 for visualization. Clustering of the filters according to filter properties can be observed.
